# Coupling of translation quality control and mRNA targeting to stress granules

**DOI:** 10.1101/2020.01.05.895342

**Authors:** Stephanie L. Moon, Tatsuya Morisaki, Timothy J. Stasevich, Roy Parker

## Abstract

Stress granules (SGs) are dynamic assemblies of non-translating RNAs and proteins that form with translation inhibition^1^. Stress granules are similar to neuronal and germ cell granules, play a role in survival during stress, and aberrant, cytotoxic SGs are implicated in neurodegeneration^2–4^. Perturbations in the ubiquitin-proteasome (UPS) system also cause neurodegeneration^5–10^, and alter the dynamicity and kinetics of SGs^11–14^. Using single mRNA imaging in live cells^15, 16^, we took an unbiased approach to determine if defects in the UPS perturb mRNA translation and partitioning into SGs during acute stress. We observe ribosomes stall on mRNAs during arsenite stress, and the release of transcripts from stalled ribosomes for their partitioning into SGs requires the activities of valosin-containing protein (VCP) and the proteasome, which is in contrast to previous work showing VCP primarily affected SG disassembly ^11, 13, 14, 17^. Moreover, members of a specialized complex in the UPS that targets aberrant nascent proteins for decay upon ribosome stalling, referred to as ribosome-associated quality control complex (RQC)^18^, are also required for mRNA release from ribosomes and partitioning into SGs. VCP alleles that increase segregase activity and cause neurodegeneration and inclusion body myopathies^5, 6, 19, 20^ increase mRNA recruitment to SGs, suggesting aberrant mRNA localization to SGs in disease contexts. This work identifies a new type of stress-activated RQC (saRQC) distinct from canonical RQC pathways in mRNA substrates, cellular context and mRNA fate.

## Main

To evaluate whether and how VCP or proteasome perturbation impacts mRNA regulation during stress, we simultaneously examined the translation status and interaction of individual mRNAs with SGs in living cells using nascent chain tracking^16^. This assay allows visualization of three distinct aspects of individual mRNA molecules being targeted to SGs including their release from polysomes, the duration of transient docking interactions with SGs, and the kinetics of stable mRNA association with SGs^15^. We examined how DBeQ, a VCP inhibitor^21^, and MG132, a proteasome inhibitor^22^ affected mRNA-nascent chain interactions and mRNA interactions with SGs in U-2 OS cells stably expressing GFP-G3BP1^23^ during acute arsenite stress, which inhibits translation initiation and leads to ribosome run-off^24^.

An important observation was that cells treated with DBeQ or MG132 during arsenite stress exhibited an increased number of mRNAs associated with nascent chains despite the presence of SGs (Fig. 1A, Movie S1, Movie S2, Movie S3). Cells were co-treated simultaneously with arsenite and either DMSO, DBeQ or MG132, and all images were taken pre-co-treatment and after stress granules formed up to 1 hour post-stress as described previously^15^. The mRNAs associated with nascent chains, in both stressed cells treated with carrier (DMSO) and in the presence of DBeQ or MG132, only interacted with SGs in a transient manner and could not enter into a stable interaction state with them (Fig. 1B), consistent with our previous observations in stressed, untreated cells^15^. A complementary approach showed a similar increase in translating mRNAs visualized with the SunTag system upon DBeQ treatment during arsenite stress^25^ (Fig. S1). These observations suggest that on some mRNA molecules VCP and proteasome function is required for efficient ribosome release during arsenite stress.

**Figure 1.**
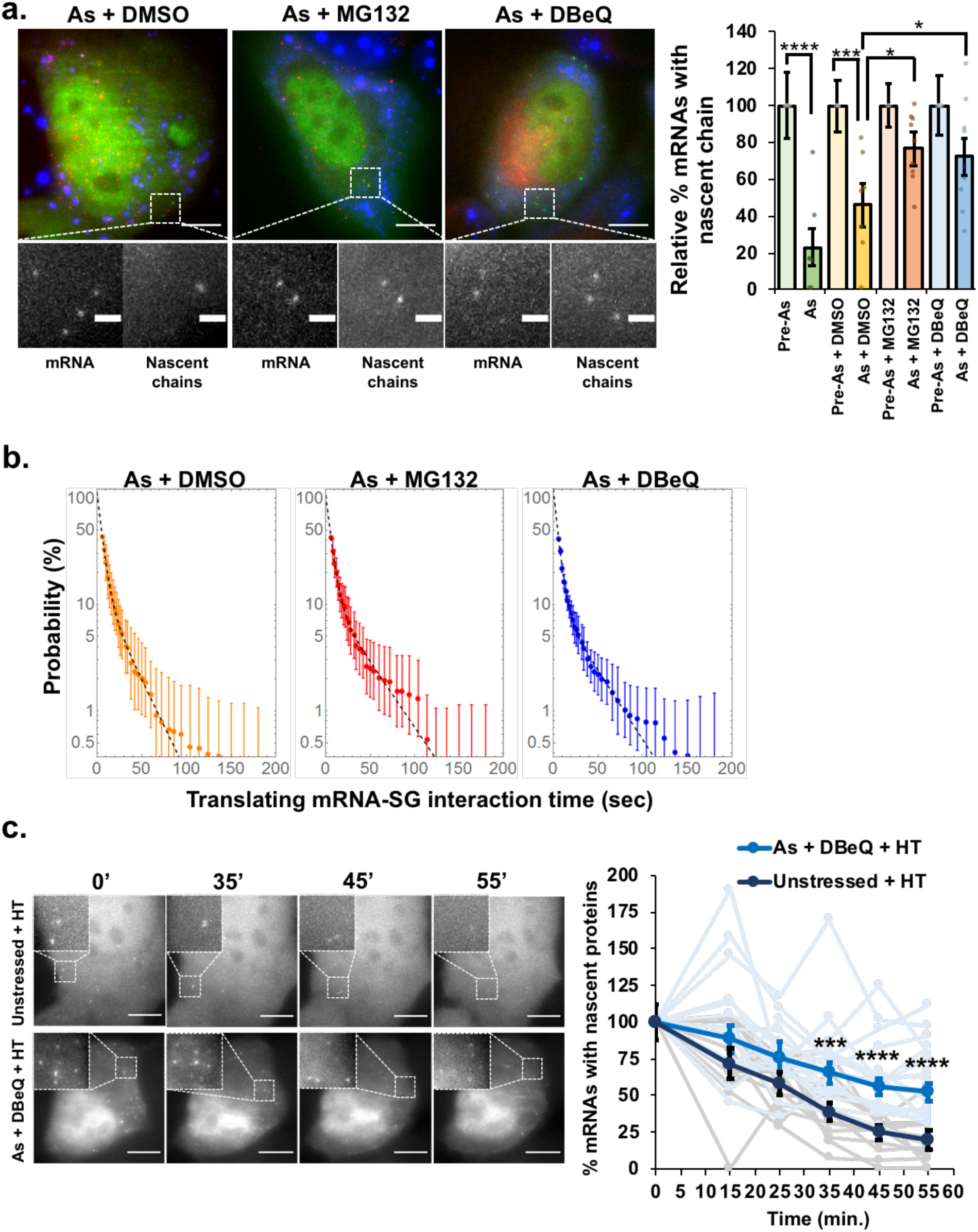
Reporter mRNAs are stalled in translation upon inhibition of VCP or the proteasome during arsenite stress. A) Left: representative images of SM-KDM5B reporter mRNAs tagged with MCP-Halo JF646 (red) with nascent chains visualized with Cy3-anti-FLAG antibody fragments (green) in cells expressing the stress granule marker GFP-G3BP1 (blue) co-treated with arsenite (“As”) and DMSO, MG132 or DBeQ (scale bars: 10 μm). Magnified panels show individual mRNAs and nascent chains (scale bars: 2 μm). Right: The average relative percent SM-KDM5B reporter mRNAs with nascent chains in individual cells +/-s.e.m. relative to pre-stress levels for As (0.5 mM, n = 7 cells), As + DMSO (0.1%, n = 7 cells), As + MG132 (10 µM, n = 9 cells) and As + DBeQ (10 µM, n = 9 cells). Student’s t-test was done to assess significance pre- and post-stress and between As + DMSO and As + MG132 or As + DBeQ with * p≤0.05, **** p≤0.001. B) Average binding-time survival probability (+/-s.e.m.) of SM-KDM5B mRNAs associated with nascent chains and SGs from n = 6 cells (As + DMSO), n = 14 cells (As + DBeQ), and n = 10 cells (As + MG132). C) Cells harboring SM-KDM5B reporter mRNAs, FLAG antibody fragments and MCP-Halo JF646 were either unstressed and treated with harringtonine (“HT”) or stressed with As and co-treated with DBeQ and HT. Images were acquired pre-stress, then in 10-minute intervals starting 15 minutes later. Harringtonine was added 15 minutes after As + DBeQ (or no treatment). Left: representative time series of unstressed (top) and As + DBeQ (bottom) cells treated with HT, scale bars: 10 μm. Right: The average +/-s.e.m. of the percent SM-KDM5B mRNAs with nascent chains at each time relative to pre-stress conditions is shown in unstressed cells (n = 13 cells, black) and stressed cells treated with DBeQ (n = 17 cells, blue), with individual cells shown in gray (unstressed + HT) and light blue (arsenite stress + DBeQ + HT). Images are maximum intensity projections from z-stacks acquired by semi-TIRF microscopy at 100x magnification. Student’s t-test was done to assess significance between untreated + HT and As + DBeQ + HT conditions at each time point with *** p≤0.005 and **** p≤0.001.

One possibility is that DBeQ and MG132 block translation repression during stress, resulting in the observed increase in nascent polypeptides associated with mRNAs. However, polysome profiling and incorporation of S^35^ -labeled met and cys into proteins revealed polysomes collapse, and translation activity is similarly suppressed during arsenite stress in the presence or absence of DBeQ or MG132 (Fig. S2) indicating co-treatments do not hyper-suppress translation. Since polysomes collapse, we infer that only a sub-fraction of mRNA molecules require VCP and proteasome function for efficient ribosome release. Therefore, VCP and proteasome inhibition do not block global translation shut-off during arsenite stress.

An alternative possibility is that those mRNAs associated with nascent chains in stressed cells treated with DBeQ or MG132 are stalled in translation elongation during arsenite stress, and ribosome run-off is inhibited. Ribosome run-off can be measured by blocking translation initiation with harringtonine and measuring the time it takes for nascent peptides chains to dissociate from the mRNA. Co-treatment of cells with the translation initiation inhibitor harringtonine revealed a significant increase in the number of mRNAs associated with nascent chains in stressed, DBeQ-treated cells compared to cells treated with harringtonine alone (Fig. 1C, Movie S4, Movie S5). However, the number of translating mRNAs decreased over time in cells co-treated with arsenite, DBeQ and harringtonine (Fig. 1C, Movie S4, Movie S5), suggesting ribosomes may be slowed rather than stopped during elongation in this context, or there are additional mechanisms for stalled ribosome removal. These results argue that ribosomes pause or are substantially slowed on a subset of mRNA molecules during arsenite stress, and require VCP and the proteasome for mRNA release.

If mRNAs are trapped in stalled translation complexes during stress upon VCP or proteasome inhibition, then recruitment of at least some endogenous mRNAs to SGs should be reduced by VCP or proteasome inhibition. By examining the SG recruitment of the *AHNAK* mRNA using single molecule fluorescence *in situ* hybridization (smFISH)^26^, we observed that the recruitment of *AHNAK* mRNAs to SGs was significantly reduced in the presence of MG132 or DBeQ during arsenite stress (Fig. 2A).

**Figure 2.**
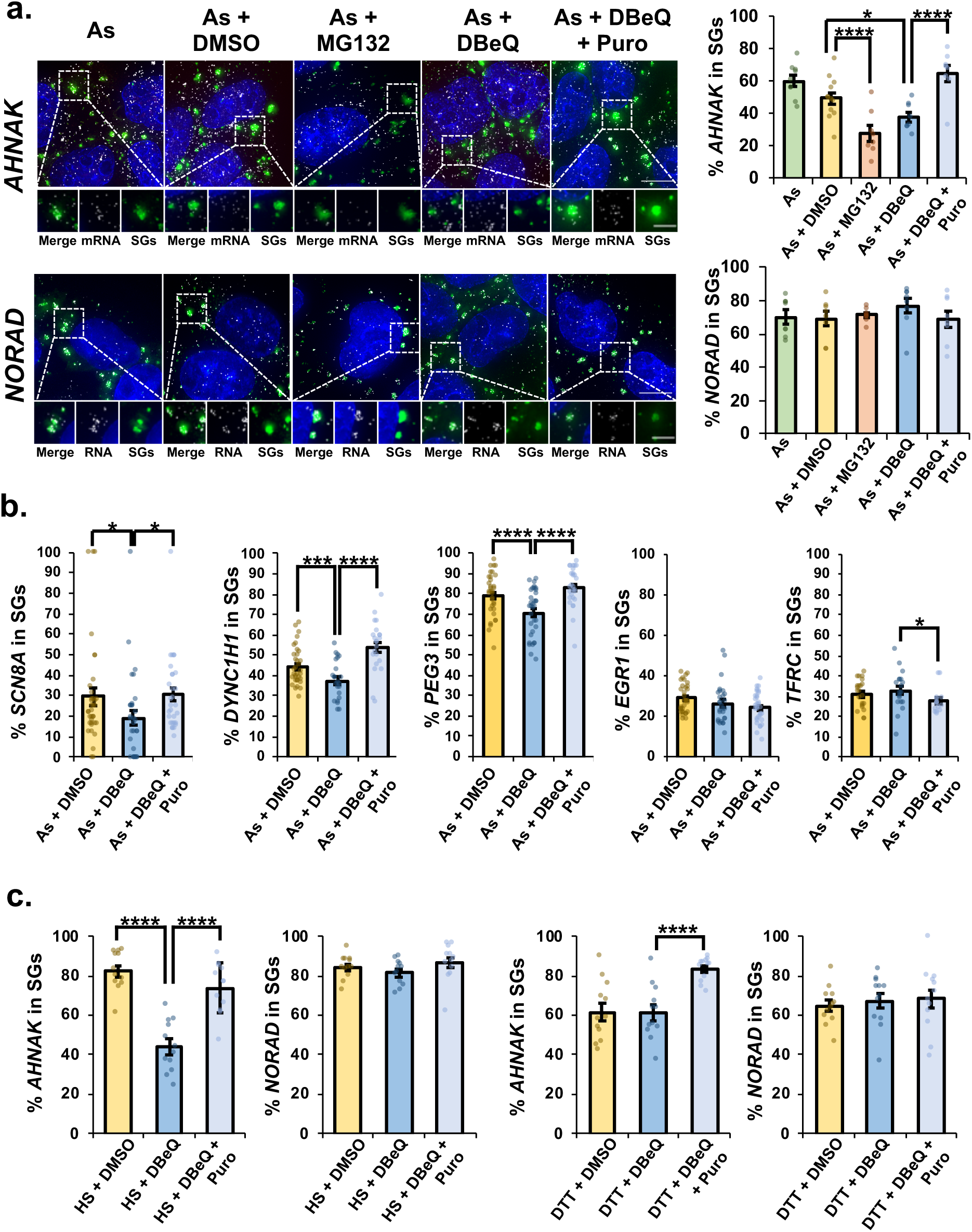
VCP and the proteasome are required for efficient release of endogenous mRNAs from ribosomes for partitioning into stress granules. **A)** U-2 OS cells expressing the SG marker GFP-G3BP1 were stressed with 0.5 mM sodium arsenite (“As”) for 45 min. with or without 0.1% DMSO (“As + DMSO”), 10 µM MG132 (“As + MG132), 10 µM DBeQ (“As + DBeQ) or 10 µM DBeQ + 10 µg/mL puromycin (“As + DBeQ + Puro”). Cells were fixed, smFISH performed to detect the mRNA *AHNAK* or lncRNA *NORAD*, and maximum intensity projections assembled from z-stacks imaged at 100x on a Delta Vision microscope. Stress granules are shown in green, RNAs in white, and nuclei in blue (DAPI). Representative photomicrographs with the average percent *AHNAK* or *NORAD* RNAs colocalizing with SGs +/-s.e.m. shown (right). For *AHNAK*, As: n = 9 cells; As + DMSO: n = 12 cells; As + MG132: n = 8 cells; As + DBeQ: n = 8 cells; As + DBeQ + Puro: n = 8 cells. For *NORAD*, As: n = 8 cells; As + DMSO: n = 8 cells; As + MG132: n = 8 cells; As + DBeQ: n = 8 cells; As + DBeQ + Puro: n = 8 cells. Two independent experiments were performed and Student’s t-test was done to assess significance, with * p≤0.05; **** p≤0.001. Scale bars: 10 µm (whole cell) or 5 µm (magnified panels). B) The average percent *SCN8A* (As + DMSO n = 36 cells, As + DBeQ n = 32 cells, As + DBeQ + Puro n = 30 cells), *DYNC1H1* (As + DMSO n = 34 cells, As + DBeQ n = 23 cells, As + DBeQ + Puro n = 22 cells), *PEG3* (As + DMSO n = 34 cells, As + DBeQ n = 34 cells, As + DBeQ + Puro n = 28 cells), *EGR1* (As + DMSO n = 33 cells, As + DBeQ n = 28 cells, As + DBeQ + Puro n = 32 cells), and *TFRC* (As + DMSO n = 22 cells, As + DBeQ n = 19 cells, As + DBeQ + Puro n = 15 cells) mRNAs in SGs +/-s.e.m. were determined as in (A) with * p≤0.05, *** p ≤0.005, and **** p≤0.001. C) The average percent *AHNAK* or *NORAD* RNAs in SGs +/-s.e.m. were determined as in (A) following heat stress (“HS”, 42°C for 45 min.), or dithiothreitol ER stress (“DTT”, 2 mM for 45 min.). Results of two independent replicates shown for HS (*AHNAK*: n = 12 cells HS + DMSO; n = 12 cells HS + DBeQ; n = 12 cells HS + DBeQ + Puro; *NORAD*: n = 12 cells HS + DMSO; n = 12 cells HS + DBeQ; n = 12 cells HS + DBeQ + Puro) and DTT stress (*AHNAK*: n = 12 cells DTT + DMSO; n = 12 cells DTT + DBeQ; n = 12 cells DTT + DBeQ + Puro; *NORAD*: n = 12 cells DTT + DMSO; n = 12 cells DTT + DBeQ; n = 12 cells DTT + DBeQ + Puro). Student’s t-test was done to assess significance, with **** p≤0.001.

Two observations suggest that *AHNAK* mRNA is limited in its recruitment to SGs due to ribosome association. First, treatment with puromycin, which releases ribosomes from mRNAs, restored targeting of *AHNAK* mRNAs to SGs (Fig. 2A). Second, the accumulation of the non-translating long non-coding RNA (lncRNA) *NORAD* into SGs was unaffected by VCP inhibition (Fig. 2A). Analyzed cells had similar numbers and sizes of SGs regardless of treatment conditions (Fig. S3).

Three other mRNAs, *SCN8A*, *DYNC1H1*, and *PEG3*, were significantly depleted from SGs in the presence of DBeQ, by ∼35%, ∼16% and ∼11% (respectively) and their localization to SGs was restored with puromycin co-treatment (Fig. 2B). However, not all mRNAs were depleted from SGs upon VCP inhibition (Fig. 2B). These observations argue ribosomes stall in translation during arsenite stress, and some, but not all, mRNAs require VCP and proteasome activity for efficient ribosome release and targeting to SGs.

To address whether VCP impacts mRNA recruitment to SGs during other stresses, we assessed the localization of *AHNAK* and *NORAD* RNAs to SGs upon heat stress and ER stress. The recruitment of *AHNAK* to SGs was reduced by ∼47% in cells exposed to heat stress in the presence of DBeQ and restored by puromycin co-treatment, while *NORAD* was unaffected (Fig. 2C). However, not all stressors cause ribosome pausing, as co-treatment with the ER stressor dithiothreitol (DTT) and DBeQ did not result in a significant reduction in *AHNAK* recruitment to SGs (Fig. 2C).

Ribosome footprinting assays have revealed ribosomes stall in mammalian cells during severe heat stress^27^ and in hyperosmotic and oxidative stresses in yeast^28^. Our results suggest that downstream of ribosome stalling, a subset of mRNAs are released from ribosomes by the action of VCP and the proteasome in acute stress contexts to target those mRNAs to SGs. One possibility is that defects in nascent chain folding are triggering this stress-activated ribosome pausing as reported in other stress contexts^27, 29^, however, ribosome or mRNA modifications may also trigger the saRQC.

VCP is known to function in the ribosome quality control (RQC) pathway where it is thought to extract ubiquitinated nascent chains from free 60S subunits after ribosome splitting. To determine if ribosome release and targeting of mRNAs to SGs involved other components of the RQC pathway, we examined if LTN1 (listerin), which ubiquitinates aberrant nascent chains, and its co-factor NEMF (nuclear export mediator factor)^18, 30^ affected mRNA targeting to SGs. We observed siRNA depletion of LTN1 (to 16.7 +/-2.4% mRNA levels), or NEMF (to 32.2 +/-4.1% mRNA levels) reduced *AHNAK* recruitment to SGs, which was restored by puromycin co-treatment (Fig. 3A). VCP, LTN1 and NEMF were not enriched in SGs and were diffusely localized in the cytoplasm with two ribosomal proteins (RPL19 and RPL29) 45 minutes post-arsenite stress in the presence or absence of DBeQ (Fig. S4), supporting a role for their function outside of SGs. The observation that VCP did not partition into SGs was in contrast to prior studies^11, 14^, however, it is possible that VCP is only recruited to SGs during late-stage stress or recovery from stress. In sum, these results point to a new function of RQC factors during stress in releasing mRNAs stalled in translation during acute stress prior to their localization to SGs.

**Figure 3.**
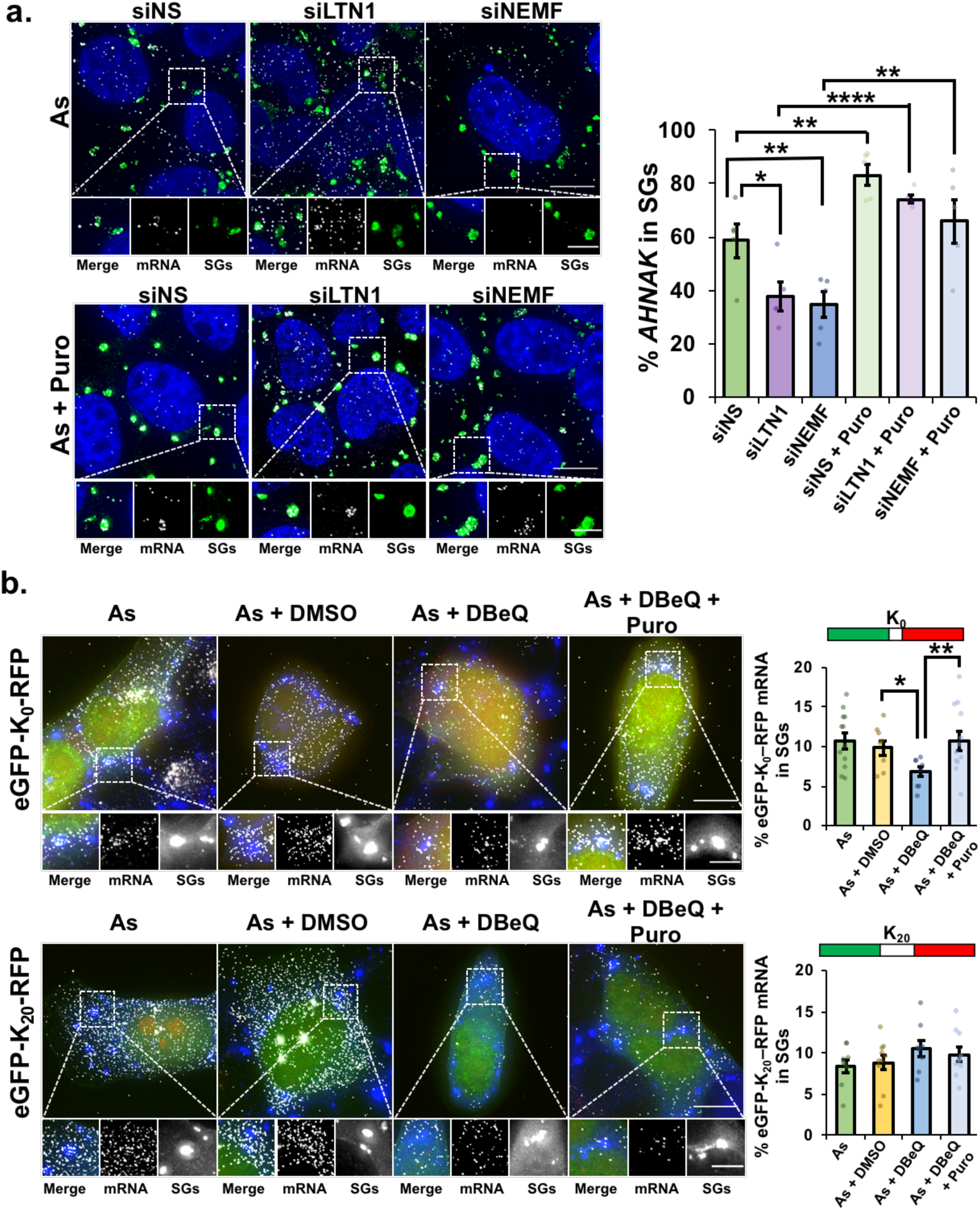
Canonical RQC factors are required for efficient mRNA partitioning into SGs, but recruitment of a canonical RQC substrate mRNA to SGs is unaffected by VCP inhibition. A) U-2 OS cells were transfected with non-specific siRNAs (“siNS”), siRNAs to *LTN1* (“siLTN1”) or *NEMF* (“siNEMF”). Cells were stressed for 45 min. with arsenite (0.5 mM) in the presence (“As + Puro”, bottom panels) or absence (“As”, top panels) of puromycin. Immunofluorescence staining to detect G3BP (green) and smFISH to detect *AHNAK* mRNA (white) was performed and nuclei visualized with DAPI (blue), representative photomicrographs at Left. Scale bars: 10 µm (whole cell) or 5 µm (magnified panels). Right: the avg. percent *AHNAK* mRNA in SGs +/-s.e.m. is shown (n = 5 frames from two independent experiments; siNS, As: 57 cells, As + Puro: 42 cells; siLTN1 As: 46 cells, As + Puro: 43 cells; siNEMF, As: 49 cells, As + Puro: 49 cells). Student’s t-test was done to evaluate significance between “siNS” and “siLTN1” or “siNEMF” conditions, and between siRNA treatments or siRNA + puromycin treatments, with * indicating p≤0.05, ** p≤0.01, **** p≤0.001. B) Dicistronic reporter mRNAs encoding eGFP and RFP separated by a linker region that encodes twenty lysine residues in a 60 nt poly(A) tract (“K20”) or no lysines (“K0”) (diagrams at top of graphs at right; from ^28^) were expressed in U-2 OS cells. Cells were stressed for 45 min. with 0.5 mM arsenite (“As”) in the absence or presence of 0.1% DMSO (“As + DMSO”), 10 µM DBeQ (“As + DBeQ”) or 10 µM DBeQ and 10 µg/mL puromycin (“As + DBeQ + Puro). Cells were fixed and G3BP detected by immunofluorescence staining (blue) to mark SGs. The reporter mRNA was detected by smFISH using probes against the eGFP open reading frame (white); eGFP is shown in green and RFP in red. Cells expressing eGFP were imaged at 100x using a Delta Vision microscope and maximum intensity projections of 25 z-stacks are shown. Representative images at Left with average +/-s.e.m. of the percent eGFP mRNA that colocalized with SGs shown at Right from two independent experiments. For K0: As n = 12 frames; As + DMSO n = 8 frames; As + DBeQ n = 8 frames; As + DBeQ + Puro n = 12 frames, with 1-5 cells counted per frame. For K20: As n = 9 frames; As + DMSO n = 10 frames; As + DBeQ n = 9 frames; As + DBeQ + Puro n = 10 frames, with 1-4 cells counted per frame. Student’s t-test was done to assess significance with *p≤0.05, **p≤0.01. Scale bars: 10 µm (whole cell) or 5 µm (magnified panels).

If a canonical RQC pathway triggered by ribosome collisions were responsible for mRNA release from stalled ribosomes during stress, an RQC substrate mRNA would be predicted to be highly sensitive to VCP inhibition in its localization to SGs. We assessed the localization of a dual-fluorescence stalling reporter mRNA with GFP encoded upstream of RFP, with or without a stalling poly(A) tract, which triggers canonical RQC, inserted between them^31^. The GFP:RFP signal intensity was higher (1.8 +/-0.3) in cells expressing the stalling reporter compared to the control reporter construct (1.0 +/-0.04), confirming lower ribosome read-through past the poly(A) sequence. We observed that while a control reporter transcript was decreased in SGs by VCP inhibition, the partitioning of the stalling reporter mRNA into SGs was unaffected by VCP inhibition (Fig. 3B). Therefore, the fate of typical mRNAs stalled in translation during arsenite stress is different from that of canonical RQC substrate mRNAs.

To determine if pathogenic VCP alleles that cause neurodegeneration and inclusion body myopathies^5^ affected mRNA release from ribosomes and partitioning into SGs, we examined *AHNAK* localization in cells expressing wild-type or eGFP-tagged pathogenic VCP variants^19, 32–35^. The common VCP-R155H variant affects the N-terminal domain which mediates ubiquitin and co-factor binding, and the severe A232E variant alters the N-terminal AAA-ATPase domain D1 which mediates homo-hexamerization of VCP and catalysis^5, 36^. In-line with previous studies that demonstrated VCP-R155H and VCP-A232E have increased ubiquitin segregase activity^19, 20^, both pathogenic variants caused increased recruitment of *AHNAK* mRNAs to SGs (Fig. 4A). Partitioning of *NORAD* into SGs (Fig. 4A), and the size and number of SGs per cell were not altered (Fig. S3), suggesting global changes in SG dynamics did not account for the increased *AHNAK* localization to SGs. It was previously shown that exogenous expression of pathogenic VCP alleles caused constitutive stress granules in a subset of unstressed cells^11^. Therefore, pathogenic VCP variants can increase the targeting of some mRNAs to SGs, which would alter the SG transcriptome, and may contribute to the formation of aberrant or constitutive SGs in disease contexts.

**Figure 4.**
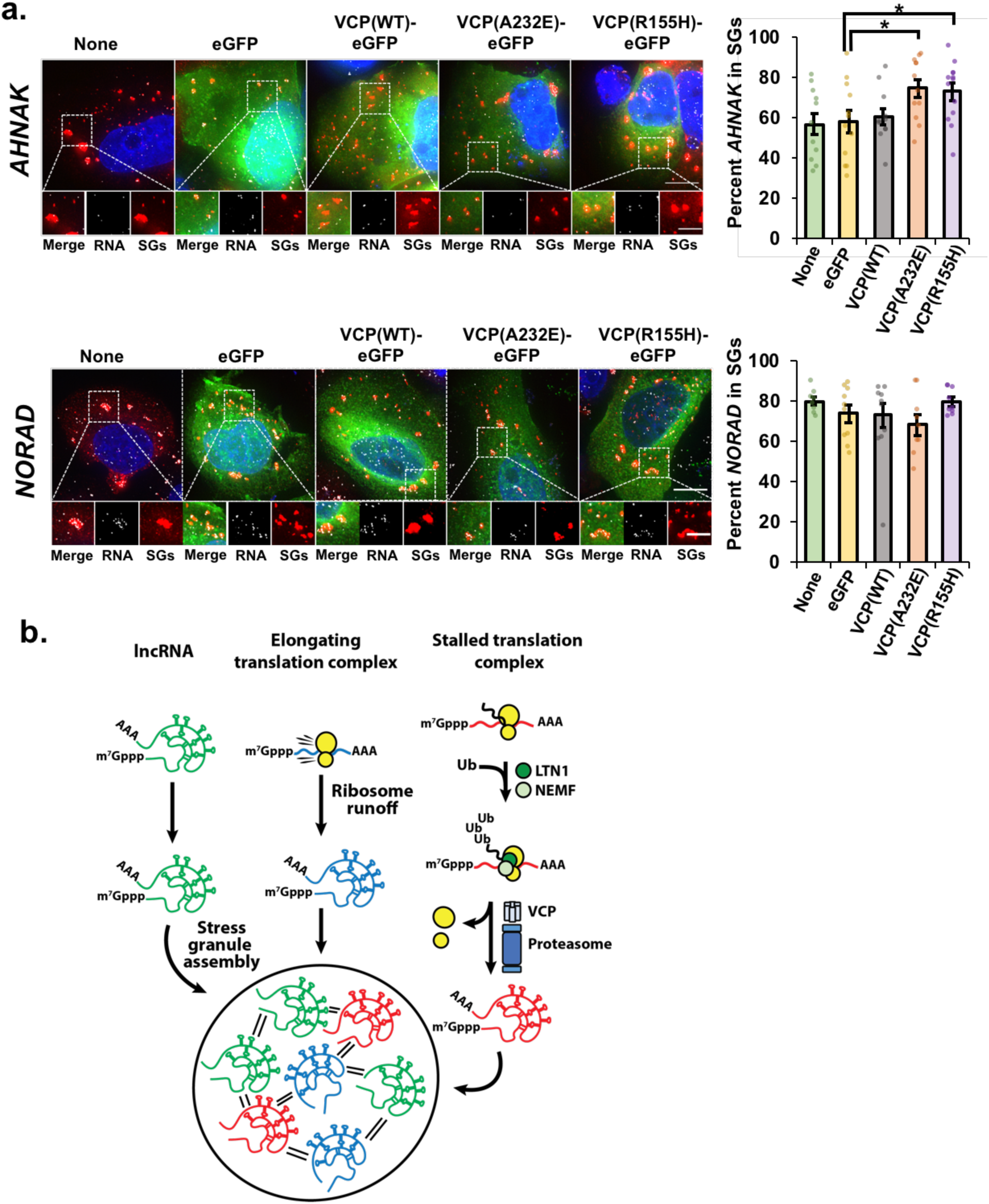
Pathogenic VCP alleles increase recruitment of the *AHNAK* mRNA, but not the lncRNA *NORAD,* to stress granules. A) U-2 OS cells stably expressing the stress granule marker mRuby2-G3BP1 were either untransfected (“None”) or transiently transfected with empty vector peGFP-N1 (“eGFP”), wild-type VCP fused to eGFP “VCP(WT)-eGFP”, or VCP alleles “VCP(A232E)-eGFP” and “VCP(R155H)-eGFP”. Cells were stressed for 45 min. with 0.5 mM arsenite then fixed, and smFISH performed to detect *AHNAK* mRNA (top) or *NORAD* lncRNA (bottom). Representative photomicrographs for are shown at left, with RNAs shown in white, stress granules in red and eGFP in green. Images were acquired on a Delta Vision microscope at 100x and maximum intensity projections of z-stacks shown. At right: avg percent *AHNAK* mRNA or *NORAD* lncRNA +/-s.e.m. that co-localize with stress granules. Three independent experiments were done and only cells expressing eGFP were counted for all transfected conditions. For *AHNAK*: None, n = 11 frames (31 cells); eGFP, n = 12 frames (25 cells); VCP(WT), n = 11 frames (17 cells); VCP(A323E), n = 12 frames (18 cells); VCP(R155H), n = 13 frames (22 cells). For *NORAD*: None, n = 8 frames (23 cells); eGFP, n = 10 frames (20 cells); VCP(WT), n = 10 frames (21 cells); VCP(A232E), n = 9 frames (12 cells); VCP(R155H), n = 9 frames (18 cells). Student’s t-test was done to assess significance between GFP and VCP(WT), VCP(A232E) or VCP(R155H) for *AHNAK* and *NORAD* with * indicating p≤0.05. Scale bars: 10 µm (whole cell) or 5 µm (magnified panels). B) Working model depicting three mechanisms by which RNAs may enter into stable associations with stress granules. Non-coding RNAs (green) can directly engage with a stress granule (large circle), while translating mRNAs exit translation via ribosome runoff (blue) or become stalled in translation (red) and must be removed by factors in the RQC complex including LTN1 (dark green circle), NEMF (light green circle), VCP (gray) and the proteasome (blue) in a process that may involve ubiquitination (“Ub”) of nascent peptides.

This study demonstrates that ribosomes stall on mRNAs during translation in acute stress conditions, and must be removed from them through a non-canonical stress-activated ribosome-associated quality control complex (saRQC) for partitioning into stress granules. We suggest a working model wherein the ubiquitin ligase LTN1 and co-factor NEMF, with established roles in nascent peptide ubiquitination in the canonical RQC^37^, trigger mRNA release from intact ribosomes by VCP and the proteasome (Fig. 4B). These experiments demonstrate the saRQC shares components of the canonical RQC pathway but diverges in molecular mechanism, mRNA substrates and mRNA fate. Specifically, in the current model of RQC, 80S ribosomes are split prior to nascent peptide release and decay from the 60S subunit, with the mRNA undergoing endonucleolytic cleavage^38^. In contrast, because we observe nascent peptides remain colocalized with mRNAs during stress, either ribosomes are not split prior to mRNA release, or there is a feedback mechanism in response to proteasome or VCP inhibition that acts upstream of ribosome splitting. In addition, saRQC and RQC act on different substrates since a traditional RQC substrate with a poly(A) tract was not altered in SG-partitioning by VCP inhibition, while a control reporter without a ribosome stall was affected. Intriguingly, VCP and the proteasome directly interact during arsenite stress^39^, suggesting the possibility that the functions and substrates of RQC factors are dynamically regulated in response to specific stress conditions.

This study suggests VCP affects mRNAs in SGs by acting on ribosome-mRNA complexes, in addition to mediating SG assembly and disassembly^11–14, 17^. One implication of this study is that SGs could serve as storage depots for mRNAs that are modified in response to stress and undergo processing by the saRQC. A second implication is that one detrimental consequence of pathogenic alleles of VCP would be increased mRNA targeting to SGs, which could alter SG properties and the cellular response to stress. Future work will aim to determine the consequences of aberrant VCP function in the saRQC on stress granule properties and function.

## Methods

### Cell culture & treatment conditions

U-2 OS and HeLa cells were grown in DMEM with 10% FBS and 1% streptomycin/penicillin at 37°C under 5% CO_2_. U-2 OS cells stably expressing GFP-G3BP1 and mRFP-DCP1a were a kind gift of Dr. N. Kedersha^23^, and those cells expressing only GFP-G3BP1 cells were isolated by flow cytometry in the BioFrontiers Institute Flow Cytometry Core facility. U-2 OS cells stably expressing mRuby2-G3BP1 (described below) were created by G418 selection and clonal expansion after plasmid transfection. HeLa cells with endogenously tagged SunTagX56-POLR2A and SunTagX32-DYNC1H1 genes stably expressing scFv-sfGFP were a generous gift of Drs. X. Pichon and E. Bertrand^25^. Cells were authenticated by morphological assessment and/or STR profiling (ATCC), and periodically confirmed negative for mycoplasma by DAPI staining. Arsenite stress was done using 0.5 mM sodium arsenite and samples were collected at 45 minutes post-stress unless otherwise indicated. For stress experiments, cells were treated with arsenite alone, DTT (2 mM) or heat stress (42°C in L-15 Leibovitz medium in CO_2_-free environment) and co-treated with DMSO (0.1%, Sigma-Aldrich), MG132 (10 μM in DMSO, Sigma-Aldrich) or DBeQ (10 μM in DMSO, Fisher Scientific) with or without puromycin (10 µg/mL, Sigma Aldrich).

### Generation of the mRuby2-G3BP1 plasmid

The mRuby2-G3BP1 plasmid was generated by insertion of codon-optimized mRuby2 lacking a stop codon into peGFP-C1-G3BP1 plasmid (a gift of Dr. N. Kedersha) to replace the eGFP coding sequence. A gBlock (Integrated DNA Technologies, shown below) was designed containing the mRuby2 sequence with 40 nt annealing arms to the peGFP-C1 G3BP1 plasmid cut with BglII and AgeI to remove the eGFP coding sequence. mRuby2 was inserted using the HiFi DNA Assembly reagent (New England Biolabs Inc).

mRuby2 gBlock sequence:

GCAGAGCTGGTTTAGTGAACCGTCAGATCCGCTAGCGCTACCGGTCGCCACCATGGTAAGCAAAGGCGAAGAACTGATTAAGGAAAACATGCGAATGAAGGTCGTAATGGAGGGCTCAGTCAATGGTCACCAGTTTAAGTGCACAGGTGAAGGTGAAGGTAGACCATACGAGGGGACACAGACCATGAGGATCAAGGTCATTGAGGGTGGCCCACTCCCATTCGCATTTGATATATTGGCGACCAGCTTTATGTACGGCTCCAGAACGTTTATTAAATATCCCGCCGATATTCCCGATTTTTTTAAGCAATCATTCCCAGAGGGATTTACTTGGGAAAGAGTCACAAGATATGAAGATGGCGGTGTCGTCACGGTTACCCAGGACACAAGTCTCGAAGATGGCGAGTTGGTTTACAACGTTAAAGTGCGGGGCGTAAATTTTCCTTCAAATGGTCCAGTGATGCAAAAGAAGACTAAAGGCTGGGAACCGAATACCGAGATGATGTACCCGGCAGACGGGGGACTGCGAGGTTACACCGATATTGCGTTGAAGGTAGATGGTGGTGGACATTTGCATTGCAACTTTGTGACCACGTATCGCAGCAAAAAGACGGTCGGAAATATAAAAATGCCTGGCGTCCACGCAGTGGACCACCGCCTCGAGCGAATTGAGGAGTCTGACAATGAAACGTATGTGGTGCAGCGCGAAGTAGCAGTGGCTAAATATTCCAATCTCGGGGGGGGAATGGACGAGTTGTATAAATCCGGACTCAGATCTATGGTGATGGAGAAGCCTAGTCCCCTGCTGGTCGG

### Nascent chain tracking

Nascent chain tracking was performed as previously described^15^ using a reporter mRNA encoding the human KDM5B ORF with 10x FLAG tags at the N-terminus and 24X MS2 stem loops inserted in the 3’ untranslated region to create the Spaghetti Monster (SM) KDM5B reporter^16^. Purified recombinant MS2 coat protein (MCP) fused with Halo and anti-FLAG antibody fragments labeled with Cy3 or Alexa 488 were bead loaded with the reporter plasmid into U-2 OS cells stably expressing GFP-G3BP1 to mark SGs. Two hours later, cells were incubated with the far-red Halo ligand JF646 for 30 minutes and washed thrice prior to imaging. Cells were imaged in complete maintenance medium lacking phenol red in a humidified chamber at 37°C under 5% CO_2_ using a custom built semi-TIRF microscope with two EM-CCD (iXon Ultra 888, Andor) cameras with a 100x oil immersion objective^16^. Cells were imaged first in the red channel, then GFP and far-red channels were imaged simultaneously. The entire cell volume was imaged with 13 z-stacks with a step size of 500 nm on a piezoelectric stage. Images were captured before stress, then were imaged every 2 sec for 10 minutes starting ∼15-45 minutes after arsenite addition (to 0.5 mM) when SGs were visible as described in^15^. For all drug treatment experiments, drugs resuspended in DMSO (10 µM DBeQ, 10 µM MG132), 0.1% DMSO or puromycin were added at the same time as arsenite.

The percent of mRNAs associated with nascent peptides was quantified in a single frame from time-lapse images obtained before and after stress using the Cell Counter plugin in ImageJ. The percent mRNAs associated with nascent peptides before stress in each cell was set to 100% and the relative percent translation after stress is presented with n = 7 cells (As), n = 7 cells (As + DMSO), n = 9 cells (As + DBeQ) and n = 9 cells (As + MG132) analyzed. One-tailed t-tests were done to assess significance between the percent mRNAs associated with nascent peptides in cells pre- and post-stress, and between cells treated with As + DMSO and As + DBeQ or As + MG132, with *p≤0.05, *** p≤0.005 and **** p≤0.001.

The duration of mRNA-SG interactions was quantified as previously described using previously published^15^ using custom *Mathematica* (version 11.2.0.0) code (accessible on GitHub at: https://raw.githubusercontent.com/TatsuyaMorisaki/Translation-Stress/master/Translation-Stress.nb) from n = 14 cells (As + DBeQ); n = 10 cells (As + MG132) and n = 6 cells (As + DMSO), from at least 2 independent experiments. The mean +/-s.e.m. binding-time survival probability of individual mRNAs associated with nascent peptides with stress granules is reported.

For harringtonine chase experiments, U-2 OS cells harboring GFP-G3BP1 or mRuby2-G3BP1 were bead loaded as above, and stressed with arsenite in the presence or absence of DBeQ (10 µM), and 15 minutes later harringtonine (3 µg/mL, Cayman Chemical Company) was added. Cells were imaged as above every 10 minutes beginning immediately prior to harringtonine addition for up to 55 minutes post-stress. The number of mRNAs associated with nascent peptides in each frame were quantified in ImageJ using the Cell Counter plugin from two independent experiments with n = 13 cells (unstressed + HT) and n = 17 cells (As + DBeQ + HT) analyzed. Student’s t-test (one-tailed) was done to assess significance between the percent mRNAs associated with nascent peptides in unstressed cells treated with HT versus cells stressed with As + DBeQ + HT at each time point. *** indicates p≤0.005, *** p≤0.001. The average +/-s.e.m. of the percent mRNAs translating at each time point is shown in bold colors (blue and black), with data from individual cells shown in light blue and gray.

### Single molecule fluorescence in situ hybridization (smFISH) and immunofluorescence (IMF) microscopy

smFISH was performed as described in^26^ using protocols adapted from Stellaris and^40–42^. Following stresses and treatments, U-2 OS cells plated on coverslips were rinsed with PBS then fixed with 10% formaldehyde or 4% paraformaldehyde in PBS for 10 minutes, washed twice with PBS, then incubated under 70% ethanol for ≥1 hr at 4°C. Cells were incubated in Wash Buffer A (10% formamide in nuclease-free 2x SSC) for 5 minutes, and hybridized with probes to *AHNAK*, *PEG3*, *TFRC*, or *NORAD* generated by Biosearch Technologies labeled in far-red (Quasar 670) or red (Quasar 570), or to *SCN8A* (Supplementary Table 3) or *EGR1* (Supplementary Table 4) generated as in ^41^, or the 5’ end of *DYNC1H1* from ^40^ in hybridization buffer (10% dextran sulfate (w/v) in 10% formamide in 2x SSC) ^42^ for 16 hr at 37°C in humidified chambers. Cells were washed with Wash Buffer A for 30 minutes at 37°C twice, then with nuclease-free 2x SSC for 5 minutes. Coverslips were mounted onto slides with VECTASHIELD® antifade mounting medium with or without DAPI (Vector Laboratories), stored at 4°C overnight and imaged as described below.

For consecutive IMF-smFISH, cells were treated and stressed as indicated, then fixed with 4% paraformaldehyde for 10 minutes. Cells were rinsed with PBS twice then permeabilized with triton-x-100 (0.1%) and RNase inhibitor (Ribolock) for 5 minutes. Cells were rinsed with PBS and incubated with mouse anti-G3BP1 (1:200, ab56574, Abcam) in PBS with RNase inhibitor (Ribolock) for 1 hour room temperature in a humidified chamber. Cells were washed thrice with PBS then incubated with goat-anti mouse Alexa-405 (1:200, A-31553, ThermoFisher Scientific) and Ribolock RNase inhibitor for SG detection for 1 hour at room temperature in a humidified chamber. Cells were washed thrice with PBS then fixed with paraformaldehyde for 10 minutes. Cells were rinsed with PBS and then the smFISH protocol described above was performed for mRNA detection. In general, SGs were detected via expression of fusion proteins GFP-G3BP1 or mRuby2-G3BP1, or via immunofluorescence detection of G3BP as indicated.

To determine protein localization (RPL19, RPL29, VCP, LTN1, NEMF and G3BP) in U-2 OS cells, and VCP and PABP localization in HeLa cells, cells were stressed and treated as indicated, rinsed with PBS then fixed with 4% paraformaldehyde for 10 minutes. Cells were washed with PBS then permeabilized with Triton-x-100 (0.5%) for 5 minutes. Cells were washed with PBS for 5 minutes then blocked in 3% bovine serum albumin (BSA) at 4°C overnight or room temperature for 1 hr. Cells were washed thrice with PBS and incubated with primary antibody (Supplementary Table 1) in 3% BSA in PBS at 4°C overnight or room temperature for 1 hr. Cells were washed thrice with PBS then incubated with secondary antibodies (Supplementary Table 1) in 3% BSA in PBS for 1 hr room temperature, then washed thrice with PBS and coverslips were mounted using VECTASHIELD antifade mounting medium with DAPI to visualize nuclei.

### Image Acquisition and Analysis of smFISH and Immunofluorescence samples

Image acquisition was performed using a DeltaVision Elite widefield microscope with a 100x oil immersion objective and PCO Edge sCMOS camera. At least 25 z-stacks were acquired (200 nm step sizes) and 2-3 frames were captured per sample in each experiment. Images were acquired and deconvolved with softWoRx (6.5.2) and maximum intensity projections created in ImageJ/Fiji^43^. For smFISH experiments, the total number of individual smFISH foci in the cytoplasm per cell, and the number of smFISH foci that co-localized with SGs were quantified using the ImageJ/Fiji^43^ Cell Counter plugin. The percent RNA in SGs was calculated for each cell or frame from 2-3 frames per replicate, and the average +/-s.e.m. from 2-3 independent experimental replicates is presented. Student’s t-test (one-tailed) was done to assess significance with *p≥0.05, **p≤0.01, ***p≤0.005 and ****p≤0.001 for all experiments. SG areas were quantified in maximum intensity projections using ImageJ/Fiji^43^ from two independent experimental replicates (two frames per condition) in cells stressed with As and stained for *AHNAK* mRNA and oligo(dT), and SGs were defined as having areas of 0.2 to >2 µm^2^ and reported (45 min. post-arsenite addition). The number of SGs per cell was quantified using the Cell Counter plugin in ImageJ/Fiji from cells stained for *AHNAK* mRNA and oligo(dT).

### siRNA-mediated depletion of LTN1 or NEMF

To deplete LTN1 or NEMF, 10 nM of siRNA SMARTpools (Dharmacon; siGENOME Human LTN1, 26046; NEMF, 9147) or a non-specific siGENOME Non-Targeting siRNA (Pool #2, Dharmacon) were transfected into U-2 OS cells with INTERFERin™ Polyplus Reagent (VWR) according to the manufacturer’s protocol. Three days later, cells were unstressed or stressed (0.5 mM arsenite) in the presence or absence of puromycin (10 µg/µL) for 45 minutes and fixed for smFISH-IMF. SGs were detected by G3BP staining, and *AHNAK* mRNA localization assessed as above.

### Transient VCP-eGFP transfections and treatments

U-2 OS cells stably expressing mRuby2-G3BP1 were plated on coverslips and untransfected or transiently transfected with peGFP-N1 (empty vector), VCP(WT)-EGFP, VCP(A232E)-EGFP or VCP(R155H)-EGFP (gifts from Nico Dantuma, Addgene #23971, #23973, and #23972)^32^. Cells were treated with 0.5 mM sodium arsenite in normal growth medium for 45 minutes at 24 hours post-transfection. Cells were fixed and smFISH performed to detect *AHNAK* or *NORAD* RNAs as described above.

Results are from three independent experiments. For *AHNAK*, a total of 31 untransfected cells (5,163 RNAs), or 25 cells expressing eGFP (2,886 RNAs), 17 cells expressing VCP(WT)-eGFP (2,089 RNAs), 18 cells expressing VCP(A232E)-eGFP (1,631 RNAs) and 22 cells expressing VCP(R155H)-eGFP (2,187 RNAs) were analyzed. For *NORAD*, a total of 23 untransfected cells (2,262 RNAs), or 20 cells expressing eGFP (2,145 RNAs), 21 cells expressing VCP(WT)-eGFP (1,957 RNAs), 12 cells expressing VCP(A232E)-eGFP (1,313 RNAs) and 18 cells expressing VCP(R155H)-eGFP (1,320 RNAs) were analyzed. Student’s t-test (one-tailed) was done to assess significance between the percent *AHNAK* or *NORAD* RNAs in SGs in cells expressing eGFP alone versus VCP(WT)-eGFP, VCP(A232E)-eGFP or VCP(R155H)-eGFP, with * indicating p≤0.05. The frequency of observing SGs with area of 0.2 µm^2^ to > 2 µm^2^ in the transfected cells analyzed for *AHNAK* localization above was obtained using ImageJ/Fiji using maximum intensity projections (histograms represent the relative frequency of each indicated SG size), and the average number of SGs per cell was derived using the Cell Counter plugin as described above.

### Metabolic labeling of nascent proteins

Nascent proteins in U-2 OS cells expressing GFP-G3BP1 were labeled with ^35^S-met and -cys (EXPRE35S35S Protein Labeling Mix, Perkin Elmer) for 30 minutes in the presence of DMSO (0.1%), DBeQ (10 µM) or MG132 (10 µM) following a 30 minute incubation in labeling medium with 10% dialyzed FBS and 1% streptomycin/penicillin. Cells were lysed in NP-40 buffer (50 mM Tris-HCl pH 8.0, 150 mM NaCl, 0.5% NP-40 substitute, 5 mM EDTA) with protease inhibitor cocktail (Sigma-Aldrich) and equal volumes were run on 4-12% NuPAGE protein gels (Thermo Fisher Scientific). Gels were exposed to phosphor screens and imaged on a Typhoon FLA 9500 phosphoimager, and ImageJ/Fiji^43^ was used to quantify the signal intensity in each lane. A representative image and the average relative nascent protein abundance +/-SEM are shown from two independent experiments. Student’s t-test (one-tailed) was done to assess significance between untreated and treated samples with ***p≤0.005. Stressed cells (0.5 mM sodium arsenite) in the presence or absence of DMSO (0.1%), DBeQ (10 µM) or MG132 (10 µM) were pulse labeled for 15 minutes starting at 0 and 15 minutes post-stress. The nascent protein abundance in each sample relative to an unstressed, untreated control was determined and individual measurements and the average +/-s.e.m. from n = 3 independent experiments are reported.

### Polysome profiles

U-2 OS cells stably expressing GFP-G3BP1 were stressed with 0.5 mM sodium arsenite or unstressed, in the presence or absence of DMSO (0.1%), DBeQ (10 µM), or MG132 (10 µM) for 30 minutes. Translation inhibitors (e.g. cycloheximide) were not used. Cells were scraped into cold PBS with protease inhibitors and RNase inhibitors (RiboLock), pelleted and frozen at −80C. Cells were resuspended in lysis buffer (500 µM Tris pH 7.5, 250 µM MgCl_2_, 150 µM KCl, 1 mM DTT, cOmplete™ ULTRA mini EDTA-free protease inhibitor cocktail (Sigma-Aldrich), RiboLock RNase inhibitor and DEPC-treated water) and vortexed. Triton-x-100 (to 0.5%) and sodium deoxycholate (to 0.5%) were added and lysates were passed through a 25G needle ten times. Lysates were clarified by brief centrifugation and total RNA was measured by A260 to load equal RNA amounts for each sample onto 15-50% sucrose gradients (15-50% sucrose in 20 mM Hepes pH 7.6, 100 mM KCl, 5 mM MgCl_2_, 1 mM DTT). Samples were ultracentrifuged at 36,000 rpm for 2 hours. Gradients were fractionated and A260 detected using a Teledyne ISCO system. The ratio of RNA in polysome and monosome fractions was determined by measuring the area under the polysome and monosome curves using ImageJ/Fiji. Two independent experiments were performed and results from one representative experiment are presented.

### RQC substrate analysis

Two constructs encoding eGFP upstream of RFP separated by a linker region encoding two P2A sites and either no lysines (“K_0_”) or 20 lysines (“K_20_”) encoded by poly(A) tracts described in ^31^ called pmGFP-P2A-K0-P2A-RFP and pmGFP-P2A-KAAA20-P2A-RFP were a gift from Ramanujan Hegde (Addgene plasmids # 105686 and # 105688; http://n2t.net/addgene:105686 and http://n2t.net/addgene:105688; RRID:Addgene_105686 and RRID:Addgene_105688). Plasmids were electroporated (BTX Harvard Apparatus Gemini System) into U-2 OS cells and 18-24 hours later, cells were unstressed or stressed (0.5 mM arsenite) in the presence or absence of DMSO (0.1%), DBeQ (10 μM) or DBeQ (10 μM) + Puromycin (10 µg/µL) for 45 minutes. Cells were fixed and immunofluorescence staining performed to detect SGs using mouse anti-G3BP (Supplementary Table 1) and goat anti-mouse 405 (Supplementary Table 1). SmFISH was performed to detect reporter mRNAs using custom made DNA probes (Integrated DNA Technologies) against eGFP designed using the LGC Biosearch Technologies Stellaris Designer (https://www.biosearchtech.com/stellaris-designer) (Supplementary Table 2). Probes were labeled via Terminal Deoxynucleotidyl Transferase (Thermo-Fisher) with ddUTP-ATTO-633 (far-red) then purified by phenol-chloroform isoamyl alcohol extraction and ethanol precipitation and resuspended to ∼12.5 µM for hybridizations (1:100). Those cells expressing eGFP were imaged and the percent eGFP mRNA in SGs was calculated as described above from two independent replicates. For K0: n = 12 frames, 5,422 mRNAs counted in As; n= 8 frames, 2,212 mRNAs counted in As + DMSO; n = 8 cells, 4,461 mRNAs counted in As + DBeQ; and n = 12 cells, with 6,672 mRNAs counted in As + DBeQ + Puro. For K20: n = 9 frames, 4,560 mRNAs counted in As; n = 10 frames, 4,500 mRNAs counted in As + DMSO; n = 9 frames, 2,964 mRNAs counted in As + DBeQ; and n = 10 frames, 1,989 mRNAs counted in As + DBeQ + Puro. Student’s t-test (one-tailed) was done to determine significance with * indicating p≤0.05 and ** p≤0.01.

### RT-qPCR

To determine the percent knockdown of LTN1 and NEMF, cells were transfected with siRNAs as above and total RNA was extracted using TRIzol^TM^ Reagent according to the manufacturer’s instructions at 72 hours post-transfection. cDNA was generated using 1 µg total RNA and random hexamers with SuperScript^TM^ III Reverse Transcriptase and qPCR performed with iQ^TM^ Supermix (Bio-Rad) using a CFX96 Real-Time Thermocycler (Bio-Rad) with *GAPDH* as a reference gene. *GAPDH* primers were previously described in^44^; *LTN1* primers (efficiency of 91.6%) that recognized both isoforms were used (LTN1 Fw: AGCAACCTGAAGCCATAGCA; LTN1 Rv: CTCTGGAACAGTTTGCGGGT), and *NEMF* primers (efficiency of 87.7%) that recognized both isoforms were used (NEMF Fw: GGTGGACGAGATCAGCAACA; NEMF Rv:

CAGCTTCAGTCAAGGTCCGT). ΔΔCq values were calculated with the Bio-Rad CFX Manager software and the average abundance of *LTN1* or *NEMF* relative to samples transfected with the non-specific control siRNAs +/-s.e.m. are reported from n = 3 independent experiments.

## Acknowledgements

We thank N. Kedersha for U-2 OS cell lines, X. Pichon & E. Bertrand for SunTag HeLa cell lines, and E. Braselmann and T. Nahreini for purifying the U-2 OS GFO-G3BP1 cell line in the BioFrontiers Institute Flow Cytometry Core facility. We thank Denise Muhlrad and Ken Lyon for technical assistance, Anne Webb and the Stasevich and Parker labs, B. Dodd and A. McCorkindale for helpful suggestions, and Jeffrey Wilusz’s lab for peGFP-N1. Spinning disc confocal microscopy was mperformed on a Nikon Ti-E microscope supported by the BioFrontiers Institute and the Howard Hughes Medical Institute at the BioFrontiers Institute Advanced Light Microscopy Core.

## Author Contributions

R.P. and S.L.M. conceptualized the study; S.L.M., T.M., R.P., and T.J.S. developed and designed methodology; T.M. and T.J.S. developed and implemented software; S.L.M. validated findings; S.L.M., T.M., and T.J.S. performed formal analysis; S.L.M. and T.M. performed experiments and/or collected data; S.L.M., R.P., T.M., and T.J.S. provided resources; S.L.M. and T.M. provided data curation; S.L.M. wrote the original draft; S.L.M., T.M., T.J.S., and R.P. reviewed and edited the drafts; S.L.M. and T.M. visualized data; T.J.S. and R.P. supervised the project; S.L.M. managed and coordinated the project; T.J.S., R.P., and S.L.M. acquired funding.

## Materials & Correspondence

Correspondence and requests for materials should be addressed to

## Code availability

Custom Mathematica (version 11.2.0.0) code previously described in Moon et al.^15^ was deposited on GitHub and accessible at https://raw.githubusercontent.com/TatsuyaMorisaki/Translation-Stress/master/Translation-Stress.nb.

## Data availability

Data supporting the findings of this study are available from the corresponding authors upon request.

## Ethics declarations Competing interests

The authors declare no competing interests.

