## Supplementary Table 1 for "Coupling of translation quality control and mRNA targeting to stress granules"

Supplementary Table 1. Antibodies used for immunofluorescence microscopy.

| Protein | Source | Dilution | Vendor | Lot number | Catalog number |
|---|---|---|---|---|---|
| **Flag (DYKDDDDK)** | Mouse | 100 μg/mL | Wako Pure Chemical Industries, Ltd | SAN4130.100 | 012-22384 |
| **LTN1 (RNF160)** | Rabbit | 1 : 100 | Novus Biologicals | A104270 | NBP1-89879 |
| **NEMF (SDCG1)** | Rabbit | 1 : 100 | Novus Biologicals | A08812 | NBP1-87924 |
| **VCP(18)** | Mouse | 1 : 50 | Santa Cruz Biotechnology | J1309 | sc-136273 |
| **RPL19** | Rabbit | 1 : 100 | Novus Biologicals | R41395 | NBP1-92346 |
| **RPL29** | Rabbit | 1 : 100 | Novus Biologicals | R99272 | NBP2-49581 |
| **PABP** | Rabbit | 1 : 500 | Abcam | GR3197139-1 | ab21060 |
| **G3BP** | Mouse | 1:200 or 1:500 | Abcam | GR3239238-1 | ab56574 |
| **Anti-mouse Alexa Fluor 594** | Goat | 1 : 1000 | Fisher Scientific | 1937185 | A-11005 |
| **Anti-rabbit Alexa Fluor 488** | Donkey | 1 : 1000 | Abcam | GR314991-3 | ab150073 |
| **Anti-rabbit Alexa Fluor 594** | Donkey | 1 : 1000 | Jackson ImmunoResea rch Laboratories, Inc. | 140419 | 711-585-152 |
| **Anti-mouse 405** | Goat | 1 : 200 | ThermoFisher Scientific | 1905835 | A-31553 |
