## Supplementary Table 2 for "Coupling of translation quality control and mRNA targeting to stress granules"

Supplementary Table 2. Oligonucleotides for smFISH to detect eGFP transcripts, related to Figure 3B.

| Probe name | Oligonucleotide |
|---|---|
| eGFP 1 | cggtgaacagctcctcgc |
| eGFP 2 | cagctcgaccaggatggg |
| eGFP 3 | ctgaacttgtggccgttt |
| eGFP 4 | gccggtggtgcagatgaa |
| eGFP 5 | agggtggtcacgagggtg |
| eGFP 6 | aagcactgcacgccgtag |
| eGFP 7 | atgtggtcggggtagcgg |
| eGFP 8 | tgaagaagtcgtgctgct |
| eGFP 9 | acgtagccttcgggcatg |
| eGFP 10 | agaagatggtgcgctcct |
| eGFP 11 | tcttgtagttgccgtcgt |
| eGFP 12 | tcgaacttcacctcggcg |
| eGFP 13 | tcgatgcggttcaccagg |
| eGFP 14 | tgaagtcgatgcccttca |
| eGFP 15 | caggatgttgccgtcctc |
| eGFP 16 | cgttgtggctgttgtagt |
| eGFP 17 | gcttgtcggccatgatat |
| eGFP 18 | gtcctcgatgttgtggcg |
| eGFP 19 | tagtggtcggcgagctgc |
| eGFP 20 | cgatgggggtgttctgct |
| eGFP 21 | ttgtcgggcagcagcacg |
| eGFP 22 | gactgggtgctcaggtag |
| eGFP 23 | gttggggtctttgctcag |
| eGFP 24 | accatgtgatcgcgcttc |
| eGFP 25 | cggtcacgaactccagca |
| eGFP 26 | cttgtacagctcgtccat |
