## Supplementary Table 3 for "Coupling of translation quality control and mRNA targeting to stress granules"

Supplementary Table 3. Oligonucleotides for smFISH to detect *SCN8A* transcripts, related to Figure 2B.

| Probe number | Oligonucleotide |
|---|---|
| SCN8A 1 | tcaggggtgaaaggcttgaa |
| SCN8A 2 | aacgtgtactccacattctt |
| SCN8A 3 | atgggtcccgtaaaaaggta |
| SCN8A 4 | gaagtttatgggccacacaa |
| SCN8A 5 | ttgtgtaaccatagttgggg |
| SCN8A 6 | ggctgctcgtaaagtcaatt |
| SCN8A 7 | ctctttcagttttatccagt |
| SCN8A 8 | ctccattgccataaacagtg |
| SCN8A 9 | atcactgagatctggaggtt |
| SCN8A 10 | taaagtgggcctgcatgaag |
| SCN8A 11 | tgtggtcctcatcaatgatg |
| SCN8A 12 | cacaaccttctgtgaagcag |
| SCN8A 13 | actttgtcagcatattccag |
| SCN8A 14 | ggctcttaagggtctcaaag |
| SCN8A 15 | tattgtcctcatacttaggc |
| SCN8A 16 | gacgattccttggattttgt |
| SCN8A 17 | tcaaaggcttgctgagtgac |
| SCN8A 18 | tagtgcctcaacgcaaacat |
| SCN8A 19 | ccaatcggatgactcggaat |
| SCN8A 20 | ccagatctcatagaaggtct |
| SCN8A 21 | cctctaacacttggattctc |
| SCN8A 22 | gtgttacgtggtaagcagac |
| SCN8A 23 | agccatttgggatcaaagtc |
| SCN8A 24 | acttttatctctaccccaaa |
| SCN8A 25 | gtgtgtgagtatgcgatgac |
