## Supplementary Table 4 for "Coupling of translation quality control and mRNA targeting to stress granules"

Supplementary Table 4. Oligonucleotides for smFISH to detect *EGR1* transcripts, related to Figure 2B.

| Probe num | Oligonucleotide |
|---|---|
| EGR1 1 | cgagtgaggaaaggatccga |
| EGR1 2 | aaaagcggccagtataggtg |
| EGR1 3 | aaatgggactgctgtcgttg |
| EGR1 4 | aaaatgtcagtgttcggcgt |
| EGR1 5 | ctgtggaaacaggtagtcgg |
| EGR1 6 | agacagaggggttagcgaag |
| EGR1 7 | tgagtggcaaaggccttaat |
| EGR1 8 | tggcaaactttcttccacag |
| EGR1 9 | cgcaagtggatcttggtatg |
| EGR1 10 | gacgggtaagaggtagcaac |
| EGR1 11 | ggggatggataagaggtagt |
| EGR1 12 | aaagcaggggggaacagagga |
| EGR1 13 | gggagaaaaggttgctgtca |
| EGR1 14 | tccatctgacctaagaggaa |
| EGR1 15 | aattggggaaggggaagtgg |
| EGR1 16 | agaacttggacatggctgtt |
| EGR1 17 | ttctagcattgaagggagca |
| EGR1 18 | aaatccatggcacagacact |
| EGR1 19 | agaggatcaccattggtttg |
| EGR1 20 | attgtcacagcatcatcaca |
| EGR1 21 | tttgccacatgtgagagtac |
| EGR1 22 | aggcaccaagacgtgaaact |
| EGR1 23 | acaattgcacatgtcaagcc |
| EGR1 24 | gccaaacagtcactttgttt |
| EGR1 25 | tgggcaataaagcgcattca |
