## Supplementary Information for "Coupling of translation quality control and mRNA targeting to stress granules"

### Supplementary Information Guide

### Supplementary Figure legends

**Figure S1. Nascent peptides marked with SunTags are elevated in arsenite-stressed cells upon VCP inhibition.** HeLa cells with endogenously tagged SunTagX32-DYNC1H1, "DYNC1H1-Suntag" and SunTagX56-POLR2A, "POLR2A-Suntag" genes and stably expressing scFv-sfGFP [22] were imaged every 10 min. in a humidified, 37ºC chamber on a spinning disc microscope at 40X for 1 hour following treatment with 0.5 mM As, co-treated with 0.1% DMSO or 10µM DBeQ. A representative time series is shown at left for the SunTagX32-DYNC1H1 cell line; scale bar: 10 µm. Shown at right is the average +/- s.e.m. of the relative percent translation foci in each frame from 7 independent experiments for each cell line. Student's t-test was done to assess significance between the relative number of translating foci in DMSO- versus DBeQ- treated cells at each time point, with * indicating $p \leq 0.05$; ** $p \leq 0.01$. Graph insets: data were fit with exponential trendlines to calculate the time when 50% of translation foci were present compared to pre-stress images for each replicate (DYNC1H1-Suntag: As + DBeQ $R^2 = 0.9527$, As + DMSO $R^2 = 0.9616$; POLR2A-Suntag: As + DBeQ $R^2 = 0.8405$, As + DMSO $R^2 = 0.8334$). Student's t-test used to assess significance with * $p \leq 0.05$, ** for $p \leq 0.01$.

**Figure S2. Translation activity is globally suppressed during arsenite stress when VCP or the proteasome are inhibited.** A) U-2 OS cells were unstressed or stressed with 0.5 mM sodium As and/or DMSO (0.1%), DBeQ (10 µM), or MG132 (10 µM) for 30 min. Cells were placed on ice and immediately pelleted and frozen at -80ºC, then lysed and polysome profiles were obtained following ultracentrifugation on sucrose gradients. Shown are representative profiles and fraction RNA in polysome:monosome ("P/M") from n = 2 independent experiments. B) Metabolic labeling of U-2 OS cells with $^{35}$S-met and cys for 30 min. was done in the presence or absence of DMSO (0.1%), 10 µM DBeQ or 10 µM MG132. Cells were lysed and proteins run on 4-12% gradient NuPAGE SDS PAGE gels and exposed to phosphorscreens. Total lane intensity was quantified with ImageJ. The average relative percent translation activity from n = 2 independent replicates is shown +/- s.e.m. Student's t-test was done to assess significance between untreated and treated cells with *** indicating $p \leq 0.005$. C) U-2 OS cells were metabolically labeled with $^{35}$S met and cys for 15 minutes immediately before collection following 0, 15 or 30 minutes under unstressed or stressed conditions (0.5 mM As) in the presence or absence of DMSO (0.1%), DBeQ (10 µM) or MG132 (10 µM). Translation activity was quantified relative to untreated, unstressed cells as in (B) from n = 3 independent experiments. The average +/- s.e.m. is shown for each condition with individual replicates shown in green (As), yellow (As + DMSO), blue (As + DBeQ) and red (As + MG132).

**Figure S3. Stress granules contain polyadenylated mRNAs, and SG area and number are not substantially altered when VCP or the proteasome are inhibited, or when pathogenic VCP alleles are expressed during arsenite stress.** A) U-2 OS cells stably expressing the SG marker protein GFP-G3BP1 (green) were stressed for 45 min. with 0.5 mM sodium As in the presence or absence of DMSO (0.1%), MG132 (10 μM), DBeQ (10 μM), or DBeQ (10 μM) and puromycin (10 μg/mL), then FISH performed with oligod(T)-Cy3 probes to detect polyadenylated transcripts (red). Nuclei are stained with DAPI (blue); these cells were also assessed for *AHNAK* mRNAs by smFISH (shown in Fig. 2A). Shown are representative maximum intensity projections of 25 z-stacks acquired at 100x on a Delta Vision microscope, scale bars: 10 μm (whole cell) or 5 μm (magnified panels). Results represent n = 2 independent experiments. B & C) Cells were treated as in (A) and the relative distribution of SG areas ($\mu m^2$) (B) and the number of SGs per cell (C) were determined using ImageJ/Fiji in the images used to quantify the fraction of *AHNAK* mRNAs co-localized with SGs (shown in Fig. 2). For As, n = 9 cells; As + DMSO, n = 12 cells; As + MG132, n = 8 cells; As + DBeQ, n = 8 cells; As + DBeQ + Puro, n = 8 cells. For cells expressing VCP alleles, results represent 3 independent experiments (None: n = 31 cells; eGFP:n = 25 cells; VCP(WT): n = 17 cells; VCP(A232E): n = 18 cells; VCP(R155H): n = 22 cells) with average number of SGs per cell +/- s.e.m. shown in (C).

**Figure S4. Components of the saRQC pathway and ribosomal proteins are not enriched in stress granules.** A) U-2 OS cells stably transfected with the stress granule marker GFP-G3BP1 (green) (left), or HeLa cells (right) were unstressed or stressed for 45 min. with 0.5 mM sodium As in the presence or absence of the VCP inhibitor DBeQ (10 μM). Cells were fixed and immunofluorescence staining performed to detect endogenous VCP (red); stress granules in HeLa cells were visualized with anti-PABP (green). Results represent n = 3 independent experiments. B) U-2 OS cells expressing GFP-G3BP1 were treated as in (A) and immunofluorescence staining done to detect Rpl29, Rpl19, LTN1, or NEMF (red). Nuclei are shown in blue (DAPI staining). Cells were imaged at 100x with a DeltaVision elite microscope and representative images are shown with scale bars: 10 μm. Results represent n = 2 independent experiments.

### Supplementary Tables

**Supplementary Table 1.** Antibodies used for immunofluorescence microscopy.

**Supplementary Table 2.** Oligonucleotides for smFISH probes to detect eGFP transcripts, related to Figure 3B.

**Supplementary Table 3.** Oligonucleotides for smFISH probes to detect *SCN8A* transcripts, related to Figure 2B.

**Supplementary Table 4.** Oligonucleotides for smFISH probes to detect *EGR1* transcripts, related to Figure 2B.

### Supplementary Movies

**Movie S1. Representative movie showing nascent chain tracking of SM-KDM5B mRNAs in U-2 OS cells treated with arsenite and DMSO, related to Figure 1A.** Nascent protein chains are shown in green, mRNAs (Halo-MS2 coat protein labeled with JF646) shown in red, and stress granules (GFP-G3BP1) shown in blue. Images were acquired every ~2 sec for 10 minutes. Scalebar is 10 $\mu$m.

**Movie S2. Representative movie showing nascent chain tracking of SM-KDM5B mRNAs in U-2 OS cells treated with arsenite and DBeQ, related to Figure 1A.** Nascent protein chains are shown in green, mRNAs (Halo-MS2 coat protein labeled with JF646) shown in red, and stress granules (GFP-G3BP1) shown in blue. Images were acquired every ~2 sec for 10 minutes. Scalebar is 10 $\mu$m.

**Movie S3. Representative movie showing nascent chain tracking of SM-KDM5B mRNAs in U-2 OS cells treated with arsenite and MG132, related to Figure 1A.** Nascent protein chains are shown in green, mRNAs (Halo-MS2 coat protein labeled with JF646) shown in red, and stress granules (GFP-G3BP1) shown in blue. Images were acquired every ~2 sec for 10 minutes. Scalebar is 10 $\mu$m.

**Movie S4. Representative movie showing nascent SM-KDM5B proteins (gray) in unstressed U-2 OS cells treated with harringtonine.** Frame 1 was acquired pre-treatment, then 15 minutes later images were obtained every 10 minutes up to 55 minutes post-stress. Harringtonine was added immediately after capturing the 15-minute image. Scalebar is 10 $\mu$m.

**Movie S5. Representative movie showing nascent SM-KDM5B proteins (gray) in arsenite stressed U-2 OS cells treated with harringtonine, related to Figure 1C.** Frame 1 was acquired pre-stress, then arsenite was added to stress cells and 15 minutes later, images were acquired every 10 minutes for up to 55 minutes post-arsenite addition. Harringtonine was added immediately after acquiring the image at 15 minutes post-stress. Scalebar is 10 $\mu$m.
