## Supplementary figures and images for "Coupling of translation quality control and mRNA targeting to stress granules"

### Figure S1

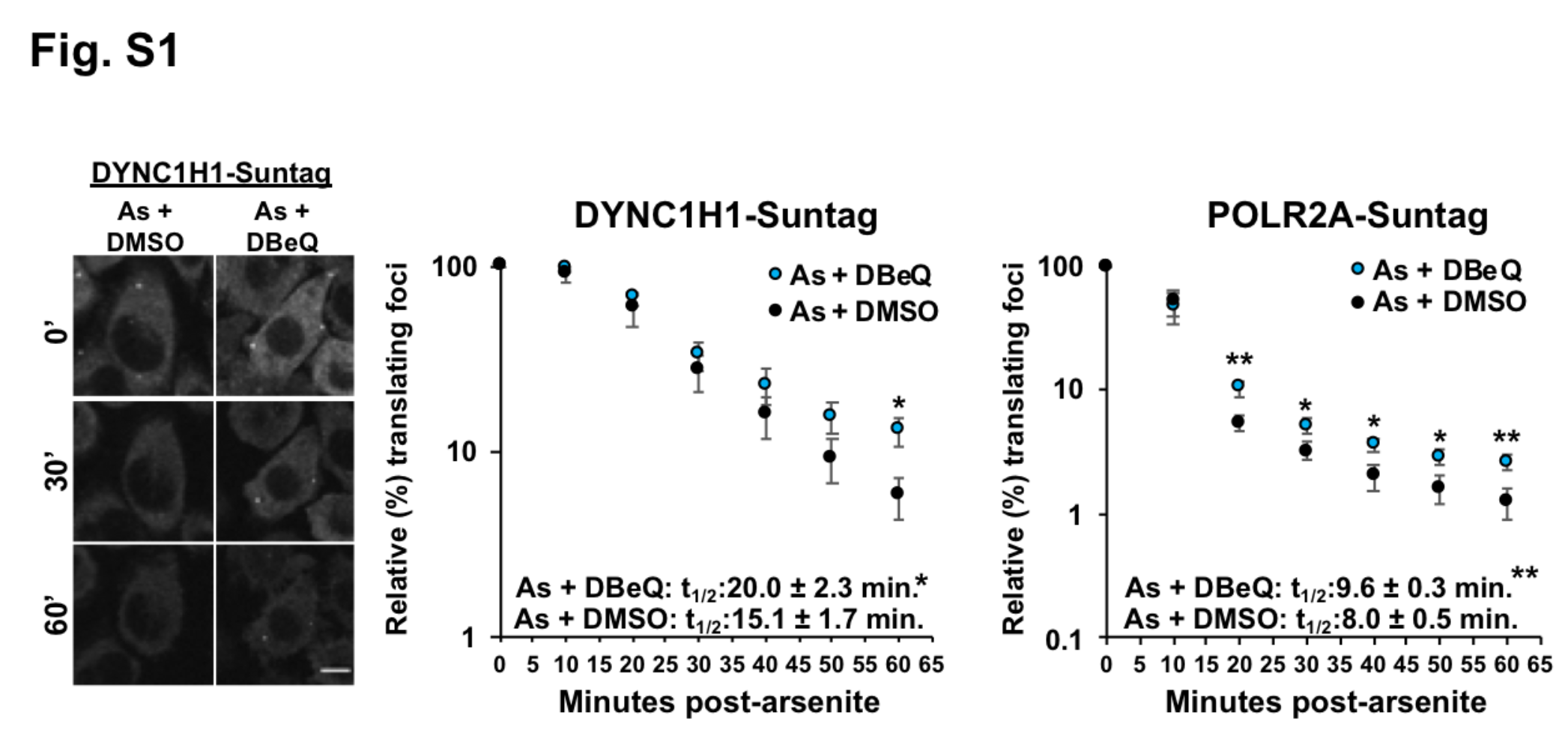

### Figure S2

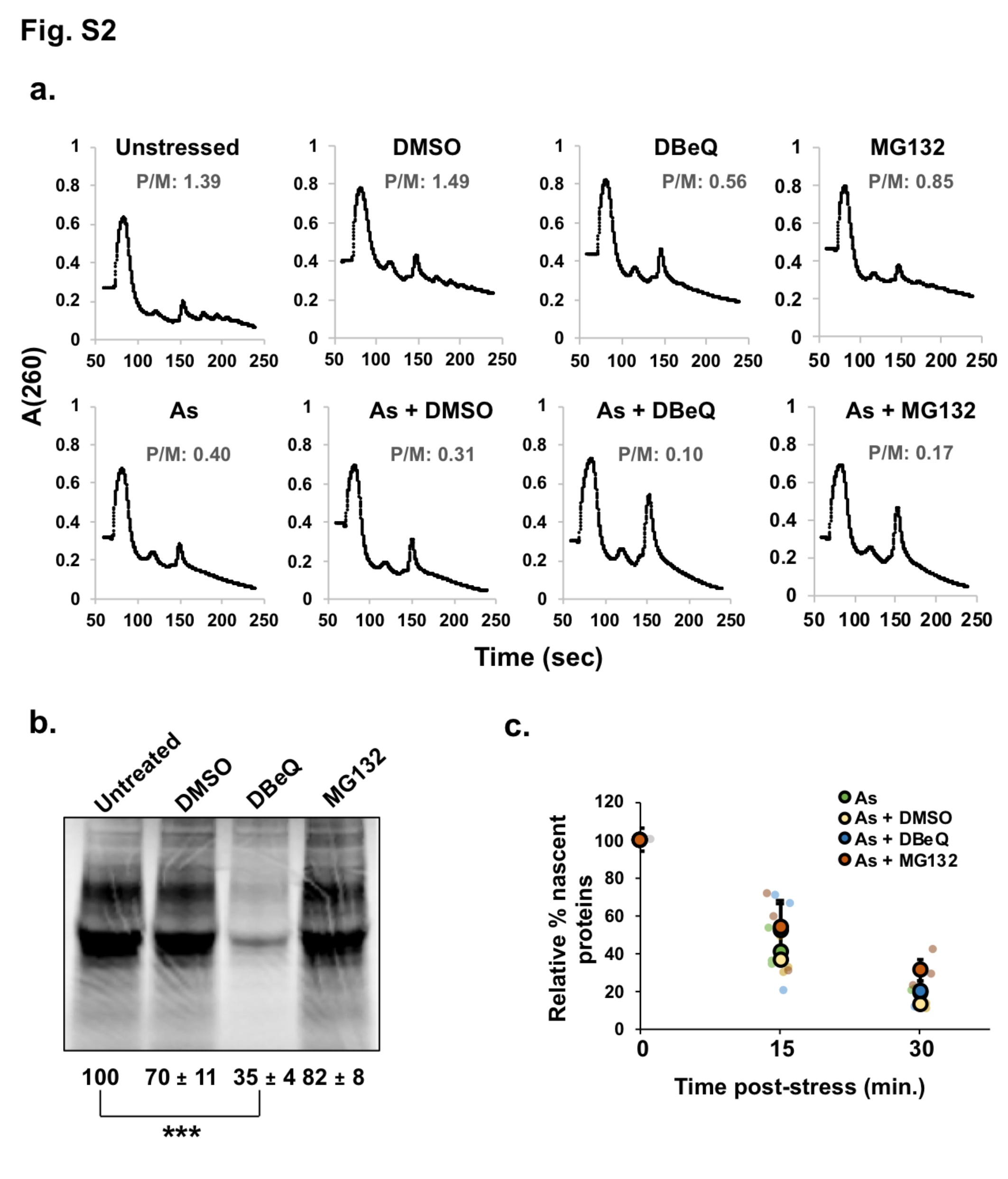

### Figure S3

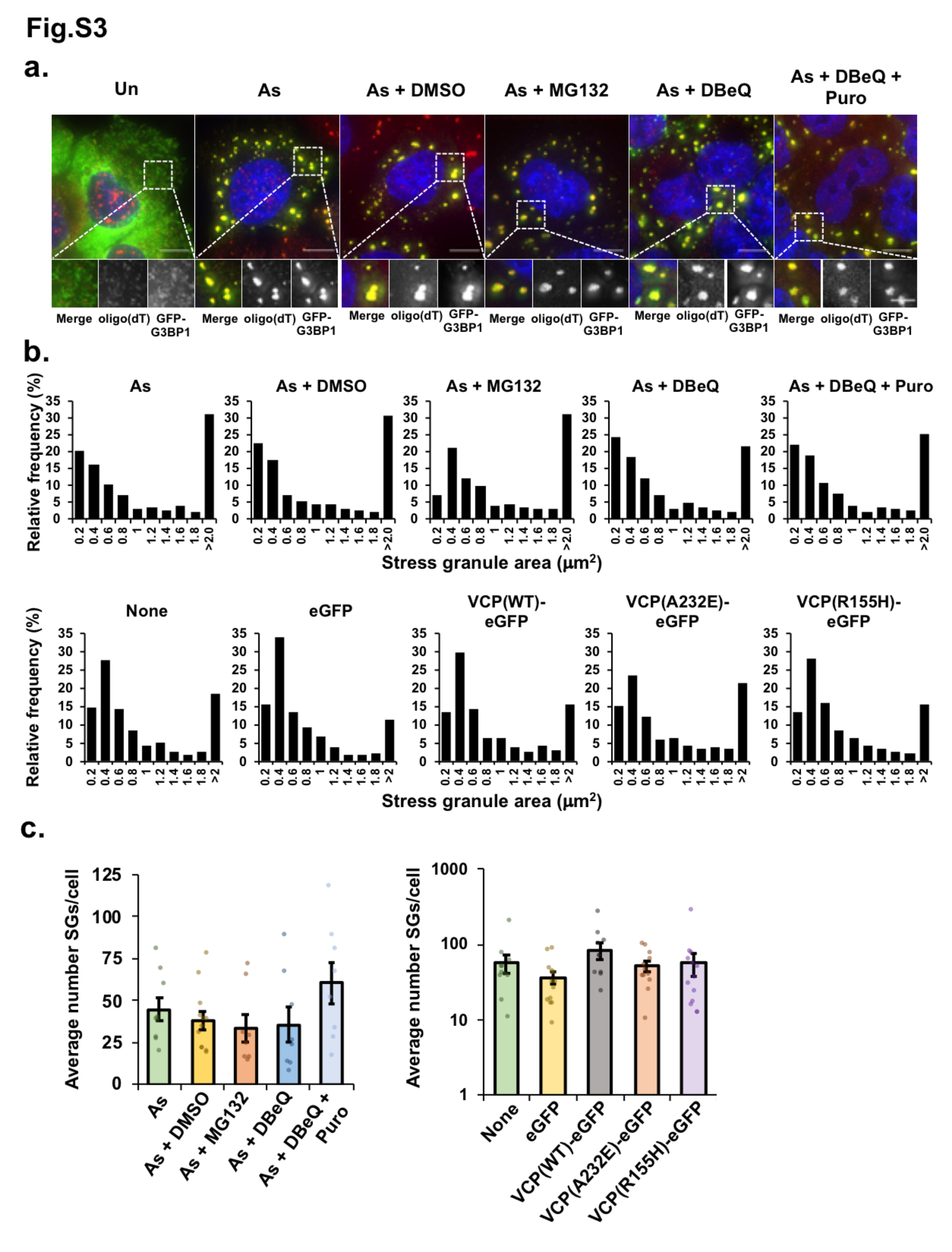

### Figure S4

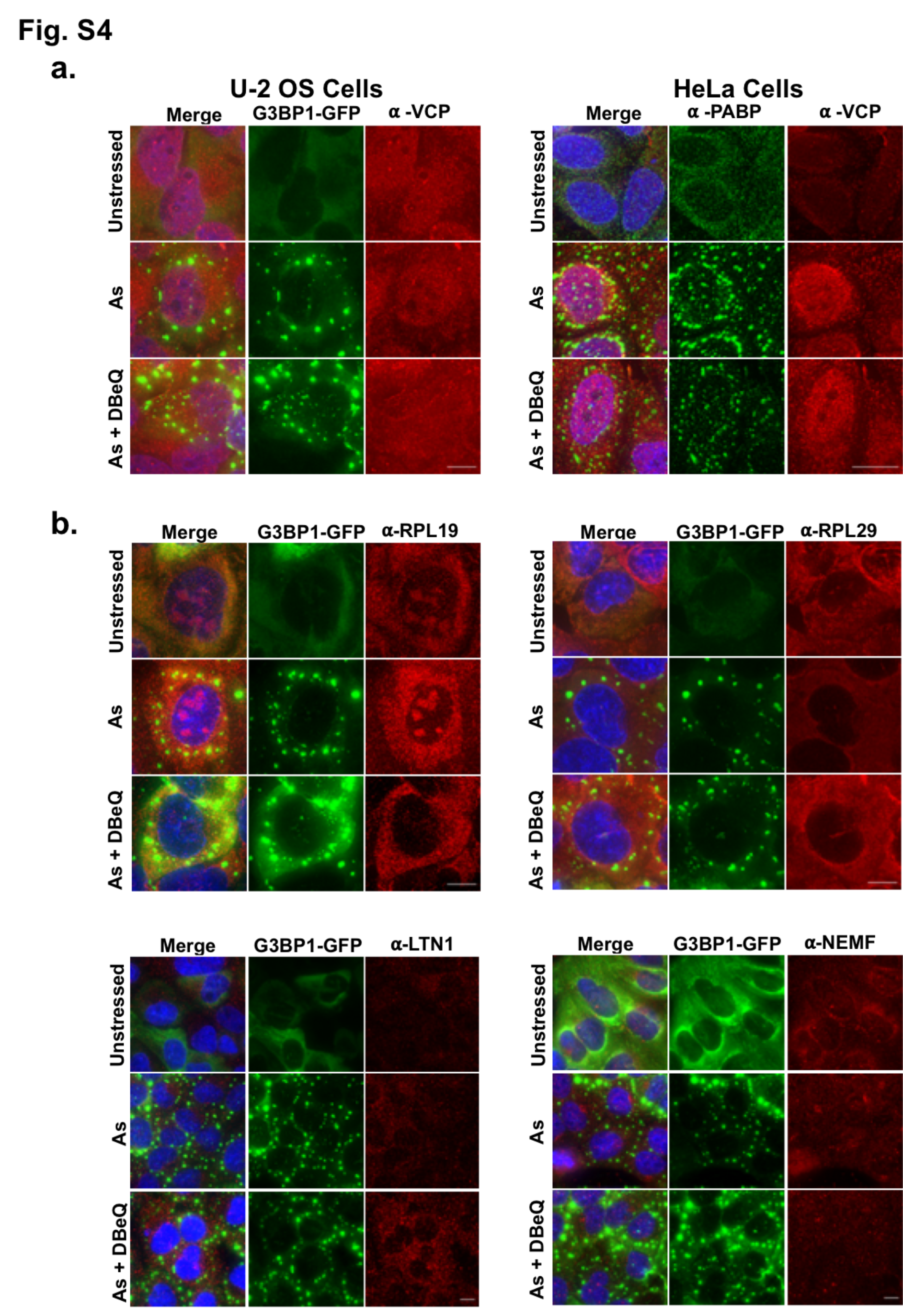
